# Investigation of adenosine A1 receptor mediated β-arrestin 2 recruitment using a split-luciferase assay

**DOI:** 10.1101/2023.02.09.527821

**Authors:** Luisa Saecker, Hanns Häberlein, Sebastian Franken

## Abstract

**Background:** Adenosine A1 receptor (A_1_AR) plays a prominent role in neurological and cardiac diseases as well as during inflammatory processes. Its endogenous ligand adenosine is known to be one of the key players in the sleep-wake cycle. Like other G protein-coupled receptors, stimulation of A_1_AR leads to the recruitment of arrestins in addition to the activation of G proteins. So far, little is known about the role of these proteins for signal transduction and regulation of A_1_AR compared to the activation of G proteins. In this work, we characterized a live cell assay for the A_1_AR mediated β-arrestin 2 recruitment. We have applied this assay to a set of different compounds that interact with this receptor.

**Methods:** Based on NanoBit^®^ technology, a protein complementation assay was developed in which the A_1_AR is coupled to the large part of the nanoluciferase (LgBiT) whereas its small part (SmBiT) is fused to the N-terminus of β-arrestin 2. Stimulation of A_1_AR results in the recruitment of β-arrestin 2 and subsequent complementation of a functional nanoluciferase. For comparison, corresponding data on the effect of receptor stimulation on intracellular cAMP levels were collected for some data sets using the GloSensor^TM^ assay.

**Results:** The assay gives highly reproducible results with a very good signal to noise ratio. Capadenoson, in contrast to adenosine, CPA or NECA, shows only partial agonism in this assay with respect to the recruitment of β-arrestin 2, whereas it is a full agonist in the case of the inhibitory effect of A_1_AR on cAMP production. By using a GRK2 inhibitor, it becomes clear that the recruitment is at least partially dependent on the phosphorylation of the receptor by this kinase. Interestingly, this was also the first time that we were able to demonstrate the A_1_AR mediated recruitment of β-arrestin 2 by stimulation with a valerian extract.

**Conclusion:** The presented assay is a useful tool for the quantitative study of the A_1_AR mediated β-arrestin 2 recruitment. It allows data collection for stimulatory, inhibitory as well as modulatory substances and is also suitable for more complex substance mixtures such as valerian extract.

## Introduction

The ubiquitous endogenous molecule adenosine is well studied and known to be part of nearly all cellular processes. It arises primarily from the breakdown of adenosine triphosphate (ATP), which is one of the major metabolites in living organisms (Sheth et al., 2014). Adenosine interacts with adenosine receptors (ARs) that belong to the superfamily of G protein-coupled receptors (GPCR). ARs are divided into four subtypes named A_1_AR, A_2A_AR, A_2B_AR and A_3_AR. They are involved in many different physiological and pathological processes and gained high interest as pharmaceutical targets (Sheth et al., 2014; Borea et al., 2018; Pasquini et al., 2022). ARs are expressed in several different cells, tissues and major organs including brain, lungs, heart, liver and kidney. A_1_AR in particular is highly expressed in the brain and central nervous system (CNS), predominantly in the cortex, hippocampus, cerebellum, spinal cord and glial cells (Fastbom et al., 1986; Reppert et al., 1991; Dixon et al., 1996). The receptors have different affinities upon adenosine, A_1_AR and A_2A_AR are considered high affine whereas A_2B_AR and A_3_AR are considered low affine (Boison, 2008). Binding of an agonist usually leads to a conformational change in the receptor resulting in activation of downstream signalling via G proteins consisting of α, β and γ subunits (Ranjan et al., 2017). A_1_AR and A_3_AR are coupled to G_i/o_ proteins resulting in inhibition of adenylate cyclase while A_2A_AR and A_2B_AR are coupled to stimulating G_s/olf_ proteins leading to stimulation of adenylate cyclase. Therefore, activation of A_1_AR and A_3_AR inhibits cyclic adenosine monophosphate (cAMP) formation with the consequence of a decreased protein kinase A (PKA) activity and phosphorylation of cyclic AMP response element binding protein (CREB). A_2A_AR and A_2B_AR vice versa increase the formation of cAMP leading to an activation of PKA and phosphorylation of CREB (Sheth et al., 2014). A_1_AR also activates phospholipase C (PLC) leading to an increase of inositol 1,4,5-triphosphate (IP3) resulting in calcium release from the endoplasmic reticulum (ER) into the cytosol (Gerwins and Fredholm, 1992). Besides G protein-dependent, ARs are also known to induce G protein-independent signalling pathways. Such pathways are initiated by receptor phosphorylation through G protein-coupled receptor kinases (GRKs) and result in binding of scaffold proteins like β-arrestins. Recruitment of β-arrestins either directs receptor desensitization or promotes downstream signalling (Klaasse et al., 2008; Shenoy and Lefkowitz, 2011; Rajagopal and Shenoy, 2018). Four different isoforms of β-arrestins are known, 1 and 4 are so called visual arrestins, 2 and 3 are non-visual. The two non-visual isoforms together with the ubiquitously expressed GRKs play a key role in the regulation of GPCR signaling (Chen & Tesmer, 2022; Gurevich & Gurevich, 2019; Shenoy & Lefkowitz, 2011). It is described that stimulated A_1_AR recruits β-arrestins. This can either lead to receptor desensitization (Klaasse et al., 2008) or mediates downstream signalling (Schulte and Fredholm, 2003; Verzijl and Ijzerman, 2011). However, there is still very limited data for A_1_AR mediated β-arrestin recruitment and signalling.

To characterize β-arrestin 2 recruitment different tools based on technologies like Förster resonance energy transfer (FRET), Bioluminescence resonance energy transfer (BRET), Tango or different kinds of protein complementation can be used (Guo et al., 2022; Hoare & Hughes, 2021; Perry-Hauser et al., 2021). In this study, a direct cellular luciferase assay using the NanoBit^®^ system was evaluated for its suitability to investigate A_1_AR ligands. The system uses a setup of two fragments that form a functional nanoluciferase when they come in close proximity. One fragment called Large BiT (LgBiT) is fused to the receptor while the corresponding smaller fragment Small BiT (SmBiT) is fused to β-arrestin 2. Luminescence as the result of the reaction catalysed by the resulting enzyme is measured in real-time. The assay system allows quantification of the specific recruitment of β-arrestin 2 initiated by compounds interacting with A_1_AR.

## Materials and Methods

### Biochemicals and reagents

HEK 293 cells were obtained from the German Collection of Microorganisms and Cell Cultures GmbH (DSMZ, ACC 305). Research reagents and chemicals were received from the following suppliers: Dulbecco’s Modified Eagle Medium (DMEM; ThermoFisher Scientific, 31885-023), Fetal Bovine Serum (FBS; ThermoFisher Scientific, A5256701), Trypsin-EDTA 0.05% (ThermoFisher Scientific, 25300062), Penicillin-Streptomycin 10.000 U/ml (PenStrep; ThermoFisher Scientific, 15140122), Phosphate Buffered Saline (PBS; ThermoFisher Scientific, 10010023), Coelenterazine h (Prolume Ltd., 50909-86-9), GloSensor^TM^ cAMP Assay Reagent (Promega, E1290), Zeocin^TM^ Selection Reagent (ThermoFisher Scientific, R25001), Hygromycin B Gold (InvivoGen, ant-hg-1), Geneticin^TM^ G-418 Sulphate (ThermoFisher Scientific, 108321-42-2). HEPES buffered Hanks balanced salt solution (HBSS/HEPES) 20 mM was freshly prepared in the lab.

The test ligands were obtained from the following suppliers: N6-Cyclopentyladenosine (CPA; Cayman Chemicals, 41552-82-3), 5’-N-Ethylcarboxamidoadenosine (NECA; Sigma-Aldrich, 35920-39-9), adenosine (Ado; Sigma-Aldrich, 58-61-7), 8-Cyclopentyl-1,3-dipropylxanthine (DPCPX; Tocris, 102146-07-6), 4-[2-[[6-amino-9-(N-ethyl-β-D-ribofuranuronamidosyl)-9H-purin-2-yl]amino] ethyl]-benzenepropanoic acid, monohydrochloride (CGS 21680; Cayman Chemicals, 124431-80-7), 2-[[6-amino-3,5-dicyano-4-[4-(cyclopropylmethoxy) phenyl]-2-pyridinyl]thio]-acetamide (BAY 60-6583; Tocris, 910487-58-0), 1-[2-chloro-6-[[(3-iodophenyl)methyl]amino]-9H-purin-9-yl]-1-deoxy-N-methyl-β-D-ribofuranuronamide (2-Chloro-IB-MECA; Cayman Chemicals, 163042-96-4), [2-Amino-4-[3-(trifluoromethyl)phenyl]-3-thienyl] phenylmethanone (VCP 171; Tocris, 1018830-99-3), 5-[2-(5-nitro-2-furanyl)ethenyl]-2-furancarboxylic acid, methyl ester (βARK1/GRK2 inhibitor; Cayman Chemicals, 24269-96-3), capadenoson (Cayman Chemicals, 544417-40-5), isopenaline hydrochloride (Iso; Sigma-Aldrich, 51-30-9), forskolin (FSK; Sigma-Aldrich, 66575-29-9). Valerian extract Ze 911 was kindly provided by Max Zeller Söhne AG (Romanshorn, Switzerland).

### Generation of expression plasmids and stably expressing cell lines

The plasmid coding for human A_1_AR fused to the N-terminus of the Large BiT (pCMV_ADORA1-LgBit) was generated by amplifying the coding region of A_1_AR by addition of a HindIII site and a BamHI site to the 5’- and 3’-end, respectively, using PCR (forward primer: 5’-GATCAAGCTTGA-TATGCCTCCCAGTATATCCG-3’; reverse primer: 5’-GATCGGATCCGATCGTCAGGCC-GTTC-3’). The plasmid ADORA1-Tango (Addgene plasmid #66209; http://n2t.net/addgene: 66209; RRID: Addgene_ 66209) used as template was a gift from Bryan Roth (Kroeze et al., 2015). The PCR product was treated with restriction enzymes and ligated via the same sites into a plasmid in front of the sequence for the LgBiT under control of the CMV promoter.

For expression of rat β-arrestin 2 with an N-terminal Small BiT (SmBit) the coding sequence was taken from pECFP-N1_rβ-arrestin-2 (a kind gift from M. Bouvier, Montreal, Canada) by restriction with NheI and SalI. The fragment was introduced into pCDNA^TM^3.1/Zeo(+) Mammalian Expression Vector (Invitrogen) containing the information for the SmBit via NheI and XhoI sites (pCDNA3.1Zeo_SmBit-β-arrestin 2).

HEK 293 cells stably expressing A_1_AR-LgBit and SmBit-β-arrestin 2 (A_1_AR-NanoBit^®^-βarr2 HEK 293 cells) were produced by double transfection using polyethylenimine (PEI) and selection of positives clones by addition of G418 (700 μg/ml) and Zeocin (100 μg/ml).

Cells used for cAMP experiments were produced by PEI-transfection of HEK 293 cells with the commercially available plasmid pGloSensor^TM^-22F cAMP (Promega, GU174434). Those cells were additionally PEI-transfected with a plasmid containing the information for the adenosine A1 receptor under control of a CMV promoter. Cells were selected using G418 (700 μg/ml) and Hygromycin B Gold (100 μg/ml; selection antibiotic of the Glosensor^TM^ system).

Stably transfected cells were cultured in DMEM low glucose supplemented with 10% (v/v) FBS, 100 IU/ml Penicillin and 100 μg/ml Streptomycin (Pen-Strep). Cells were incubated at 37°C and 5% (v/v) CO_2_. Cells were passaged every 3 days in a ratio of 1:10 at a confluency of 80-90%.

### Establishment of cell-based assay systems

#### β-arrestin 2 recruitment assay

The assay was constructed to detect and monitor real-time protein-protein interactions. Once the two fragments LgBiT and SmBiT come in close proximity after receptor activation and phosphorylation they build a functional enzyme that is generating light upon addition of its substrate fumerazine or coelenterazine (Wouters et al., 2018).

A_1_AR-NanoBit^®^-βarr2 HEK 293 cells were seeded in a white clear bottom 96-well plate (25.000 cells/well) and incubated at 37°C and 5% CO_2_ for 24 hours. Test compounds were either diluted in DMSO 10% or water; however, the maximum concentration of DMSO on cells was 1%. The medium was replaced by 45 μl Coelenterazine h substrate solution (2.5 μM Coelenterazine h in HEPES buffered Hanks balanced salt solution). The 96-well plate was immediately placed into the Tecan Spark^®^ multimode microplate reader where the background luminescence was measured 3 to 5 cycles until a stable signal was obtained. The measurement was paused and cells were stimulated with 5 μl ligand solutions. The measurement was continued for another 55 to 57 cycles.

#### GloSensor™ cAMP assay

Establishment of HEK 293 cells expressing a cAMP biosensor as well as measurement of cAMP were performed as described by Bussmann et al. (Bussmann et al., 2020). Cells were seeded in a white clear bottom 96-well plate (35.000 cells/well) and incubated at 37°C and 5% CO_2_ for 24 hours. The medium was changed to 25 μl substrate solution per well containing 4% GloSensor™ cAMP Reagent stock solution in HEPES buffered DMEM. The cells were incubated for one hour at 37°C and subsequently equilibrated for another hour at room temperature in the plate reader (Tecan Spark^®^). Cells in which adenylate cyclase had been activated with forskolin and isoprenaline were stimulated with different A_1_AR agonists. Decrease in cAMP concentration due to A_1_AR stimulation was measured by luminescence differences.

#### Data analysis and statistics

In case of the NanoBit^®^ assay, raw data of three to four independent experiment was collected and transferred to Prism (Graph-Pad Software, Inc., San Diego CA, USA; version 9.5.0) and plotted as a function of time. To normalize for well to well variabilities each data point was divided by the mean of the first three values using the “remove baseline and column math” function of the software. Next the solvent control was subtracted using the same functionality of Prism. After absolute signals were corrected for solvent control samples and inter-well variability, areas under the curves (AUC) were calculated using the corresponding function and plotted against the concentration of the agonist used (log M). A sigmoidal curve was fitted to calculate EC_50_ and IC_50_ values based on the dose-response data by using a nonlinear regression model (variable slope; see also supplemental information Figure 1).

**Figure 1:**
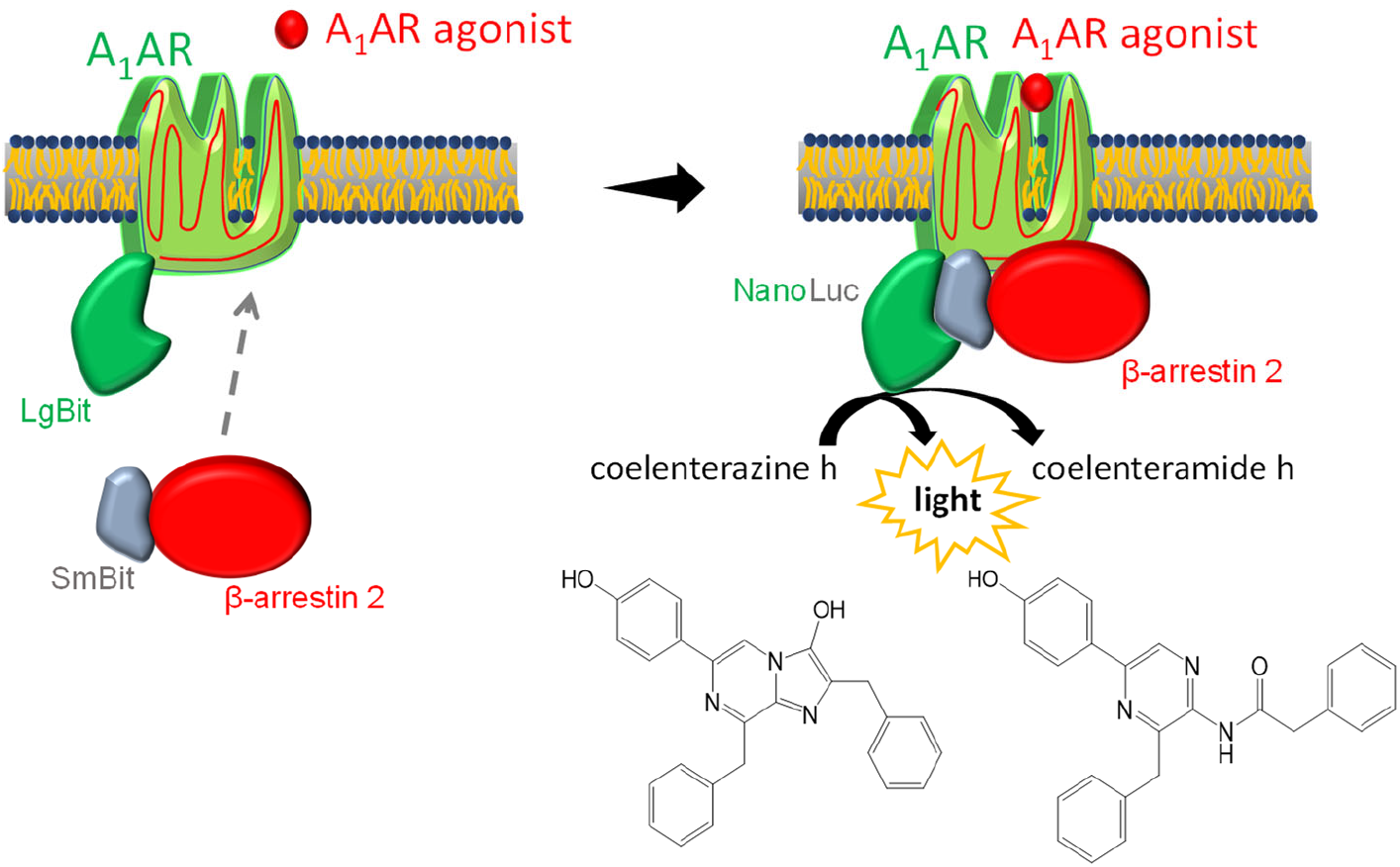
Luciferase-based β-arrestin 2 recruitment assay. After agonist binding SmBiT-β-arrestin 2 migrates to adenosine A1 receptor (A_1_AR) that is coupled to LgBiT. Binding of LgBiT and SmBiT results in a functional enzyme (NanoLuc) that generates light in the presence of the substrate coelenterazine h.

Initial rates were calculated using the plug-in equations provided by Dr. Samuel Hoare (Hoare, Tewson, Quinn, & Hughes, 2020). Specifically, the rise-and-fall equation that consider baseline and drift was used to calculate initial rates.

For Glosensor™ cAMP assay raw data of single experiments was transferred to Prism and plotted as a function of time. IC_50_ values based on the dose-response data were calculated as described above. All experiments were repeated at least twice to verify results.

To detect statistical differences between groups, analysis of variance (ANOVA) followed by post hoc analysis (Tukey) was performed.

## Results

### Different A_1_AR agonists show different efficiencies for β-arrestin 2 recruitment

The established cell-based NanoBit^®^ assay was designed to study β-arrestin 2 recruitment by adenosine A1 receptor in real time. For this purpose, the LgBiT was linked to the C-terminus of the receptor and the SmBiT to the N-terminus of β-arrestin 2 (Fig. 1). Similar to what was seen for other adenosine receptors (Storme, Cannaert, et al., 2018), this combination resulted in a dose-dependent signal increase after application of adenosine to transfected HEK 293 cells (Fig. 2 A). Using these data sets, an EC_50_ for adenosine of 884 nM could be calculated. Furthermore, dose-response experiments for CPA (Fig. 2 C) as well as NECA were performed (Fig. 2 C). As already seen for adenosine very robust dose response curves for both agonists could be calculated from the AUC values.

**Figure 2:**
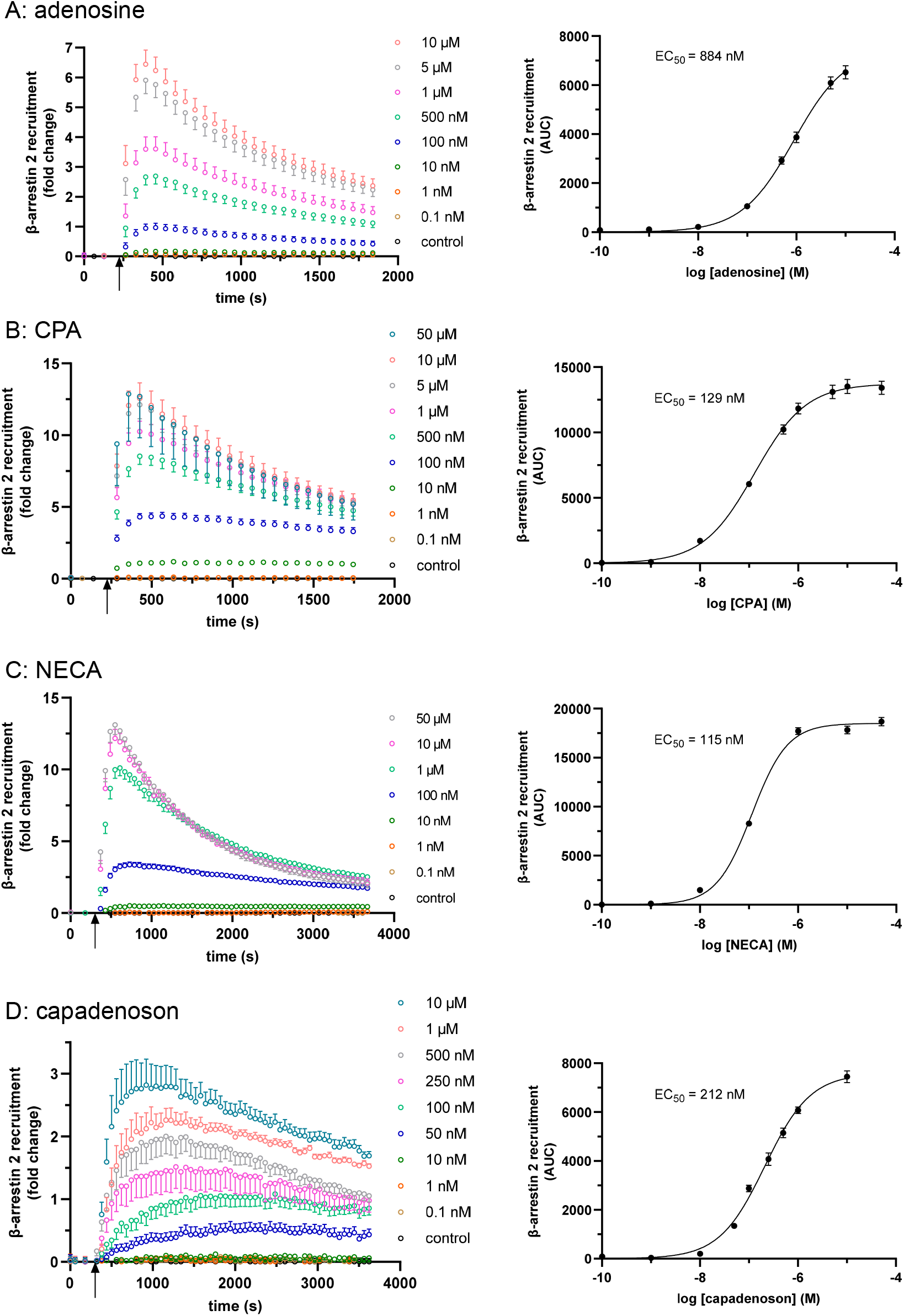
Concentration-dependency of β-arrestin 2 recruitment in the A_1_AR NanoBit^®^ reporter assay: At the time point indicated by the arrow different agonists were added to A_1_AR-NanoBit^®^-βarr2 HEK 293 cells at concentrations indicated beside the graph and luminescence was measured for up to 60 minutes. A solvent control of 0.1% DMSO was included. Dose response curves on the right were calculated using the areas under the curves (AUC). Values are given as mean +/− SEM (n=9 of three independent experiments).

Both agonists showed significantly lower EC_50_ values for β-arrestin 2 recruitment compared to adenosine: 129 nM (CPA) and 115 nM (NECA), respectively.

Next, we tested the non-nucleoside agonist capadenoson for recruitment of β-arrestin 2. It induced a dose-dependent recruitment which was less pronounced than for the other agonists tested. The calculated EC_50_ for capadenoson was 212 nM (Fig. 2 D). When tested side by side in a saturated concentration, fold change of luminescent signal was about 4-5 times higher for adenosine, CPA, and NECA than for capadenoson (Fig. 3 A). Calculating its efficacy compared to CPA capadenoson reached only 41.8% (see table in Fig. 3 C).

**Figure 3:**
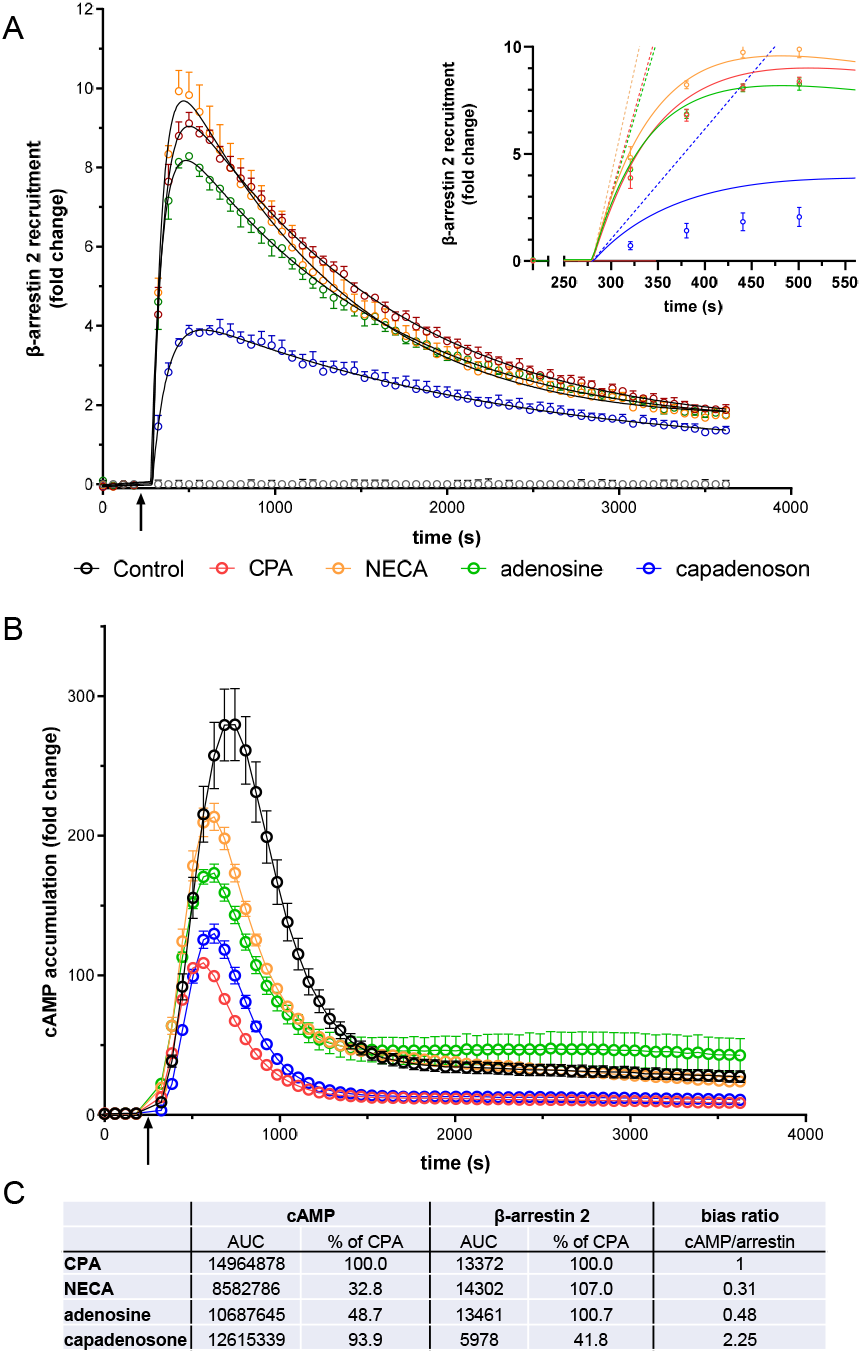
(A) β-arrestin 2 recruitment in the A_1_AR NanoBit^®^ reporter assay by different agonists: 10 μM of either CPA, NECA, adenosine or capadenoson were added to A_1_AR-NanoBit^®^-βarr2 HEK 293 cells at the time point indicated by the arrow and luminescence was measured for 60 min. A solvent control of 0.1% DMSO was included. Curves calculated by Prism plug in “rise-and-fall equation” provided by Dr. Samuel Hoare are given in black. Insert: Activation phase. Initial rates calculated from the fitted curves are given as dashed lines. Values are given as mean +/− SEM (n=12 of four independent experiments). (B) cAMP production measured by the A_1_AR-GloSensor™ assay induced by different agonists: 1 μM of either CPA, NECA, adenosine or capadenoson were added to GloSensor^TM^ HEK 293 cells at the time point indicated by the arrow and luminescence was measured for 60 min. A solvent control of 0.1% DMSO was included. Values are given as mean +/− SEM (n=9 of three independent experiments). (C) Table giving the AUC values for the different conditions in both assays, the efficacy related to CPA and the ratio between cAMP production and β-arrestin 2 recruitment.

Different specific and non-specific A_1_AR agonists show different efficacy profiles concerning cAMP formation compared to β-arrestin 2 recruitment. For comparison with the β-arrestin 2 results, we measured all agonists in a saturated concentration for cAMP production in an A_1_AR GloSensor™ cAMP assay (Fig. 3 B). All agonists tested led to a significant reduction of cAMP production with CPA showing the strongest effect. The efficacies of the substances were calculated as percentage of the efficacy of CPA (see table in Figure 3 C).

Calculation of the AUC contains activation as well as deactivation/internalization events of the receptor. In some cases, it can be useful to compare the activation of the receptor only. Hoare et al. developed a tool for evaluating G protein- and arrestin-mediated signaling in live cells, which allows calculation of receptor activation and deactivation separately (Hoare & Hughes, 2021). This tool was developed using fluorescent reporters but curves calculated by it fitted quite well to our lumines-cence-based data. Figure 3 A shows the fit for the different agonists used in this study. The analysis tool provides different parameters, such as the initial rate. This is the slope of a straight line adapted to the activation phase (see box in Figure 3 A). The initial rates calculated from the fits were 7.52 +/− 0.01286-fold change/min for adenosine, 6.95 +/− 0.01194-fold change/min for CPA, and 9.04 +/− 0.01515-fold change/min for NECA but only 1.55 +/− 0.00854-fold change/min for capadenoson.

### Specificity of A_1_AR NanoBit^®^ reporter assay for β-arrestin 2 recruitment

Further on, the specificity of the assay was investigated with the AR agonists NECA (nonspecific), adenosine (nonspecific), CPA (specific, A_1_AR), CGS 21680 (specific, A_2A_AR), BAY 60-6583 (specific, A_2B_AR), and 2-Cl-IB-MECA (specific, A_3_AR). Untreated or isoprenaline-treated cells served as controls. The agonists were tested at two concentrations, 0.1 μM and 1 μM. As seen already before, both nonspecific AR agonists NECA and adenosine as well as the specific agonist CPA caused significant recruitment of β-arrestin 2 (p < 0.01). Treatment with NECA (10.288 +/− 1065 AUC and 20.204 +/− 2917 AUC) and CPA (14.588 +/− 1785 AUC and 21.056 +/− 2144 AUC) resulted in higher recruitment compared to adenosine (5208 +/− 410 AUC and 12288 +/− 1658 AUC). In contrast, agonists specific for the other three adenosine receptors did not significantly increase β-arrestin 2 recruitment (Fig. 4 A). To further test the specificity of the assay, the effect of DPCPX, an A_1_AR inhibitor, on recruitment was examined. DPCPX blocked β-arrestin 2 recruitment by CPA in a dose-dependent manner (Fig. 4 B). The calculated IC_50_ for DPCPX was 124 nM.

**Figure 4:**
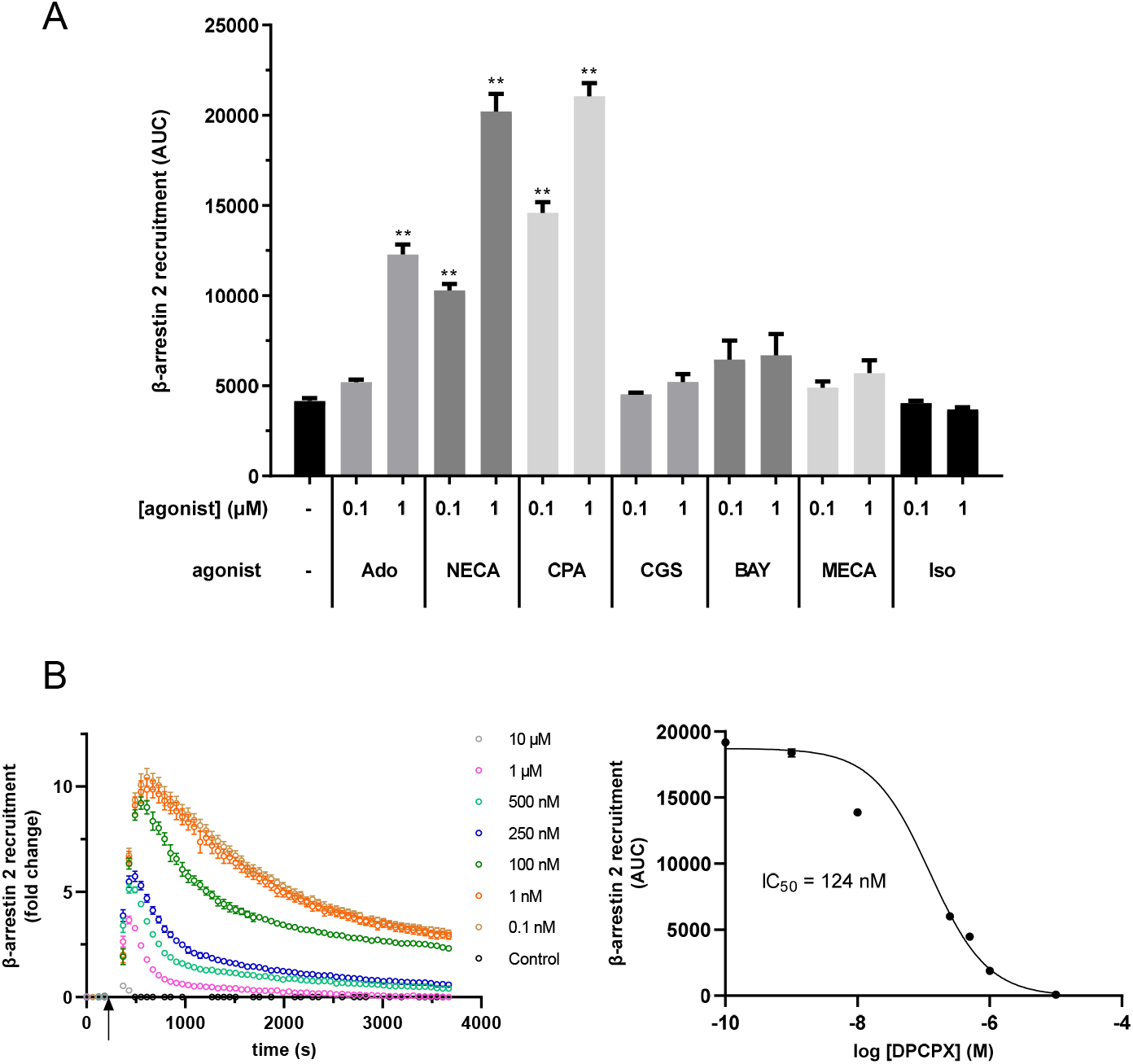
(A) Different adenosine receptor agonists were tested in the A_1_AR NanoBit^®^ reporter assay in concentrations of 0.1 and 1 μM, respectively. Adenosine (Ado; unspecific, endogenous agonist), NECA (unspecific agonist), and CPA (specific A_1_AR agonist) led to a significant increase in β-arrestin 2 recruitment whereas CGS 21680 (CGS; specific A_2A_AR agonist), BAY 60-6583 (BAY; specific A_2B_AR agonist), and 2-Cl-IB-MECA (MECA; specific A_3_AR agonist) did not changed the recruitment significantly when compared to control (0.1% DMSO). Isoprenaline (Iso; β-adrenergic agonist) was used as unrelated control. Values are given as mean +/− SEM (n=9 of three independent experiments). (B) Influence of the A_1_AR antagonist DPCPX on β-arrestin 2 recruitment in the A_1_AR NanoBit^®^ reporter assay. A_1_AR-NanoBit^®^-βarr2 HEK 293 cells were preincubated with increasing amounts of DPCPX. β-arrestin 2 recruitment was induced using 1 μM CPA. Right: Dose-response curve for DPCPX. Values are given as mean +/− SEM (n=9 of three independent experiments); **p<0.01.

### Modulation of A_1_AR/β-arrestin 2 interaction measured by NanoBit^®^ reporter assay

Beside agonistic and antagonistic effects modulation on GPCRs can occur. One such modulator of A_1_AR is VCP 171, a 5-substituted 2-aminothiophene. Co-incubation of NanoBit^®^ HEK cells with VCP 171 increased the agonistic effect of NECA (Fig. 5 A). β-arrestin 2 recruitment increased by 10%, 18%, and 19% for 0.1 nM, 1 nM, and 100 nM NECA, respectively.

**Figure 5:**
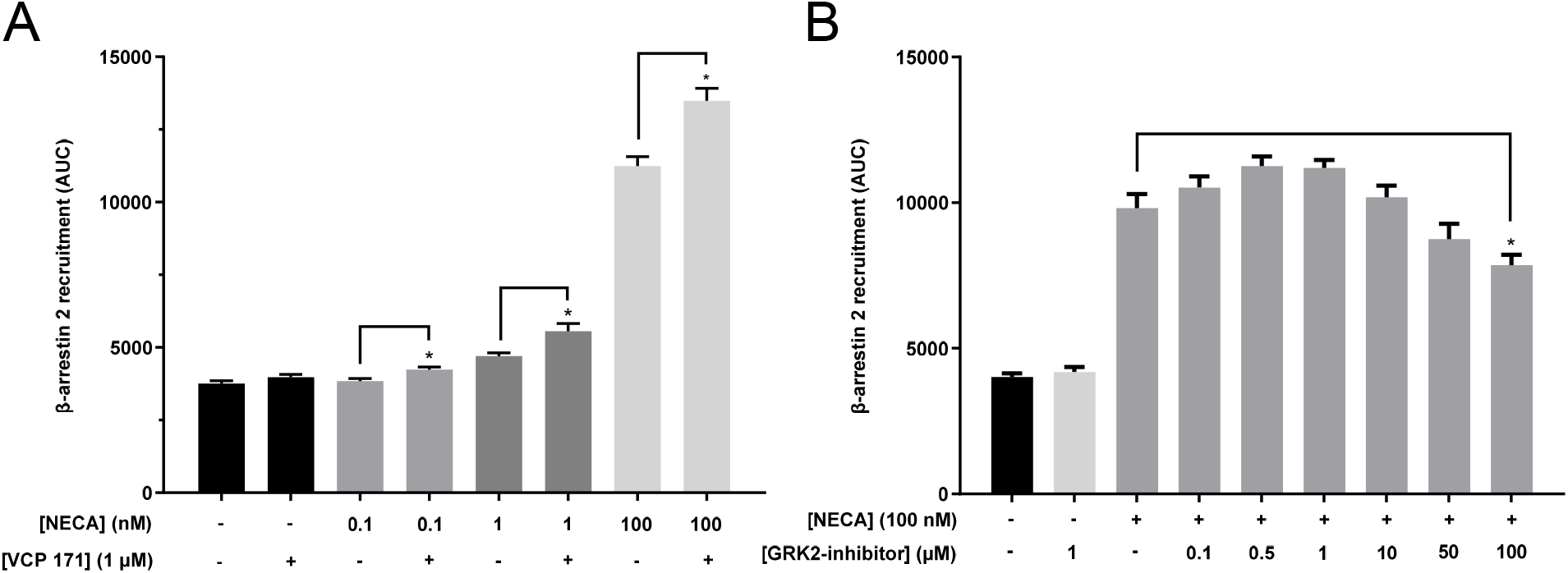
Modulation of β-arrestin 2 recruitment by the A_1_AR. (A) A_1_AR-NanoBit^®^-βarr2 HEK 293 cells were stimulated with 0.1 nM, 1 nM, or 100 nM NECA without or with 1 μm of the positive A_1_AR modulator VCP 171. The combination of VCP 171 and NECA resulted in a significant increase in β-arrestin 2 recruitment when compared to cells treated with NECA alone. (B) A_1_AR-NanoBit^®^-βarr2 HEK 293 cells were stimulated with 100 nM NECA in the presence of different concentrations of GRK2 inhibitor (βARK1). Incubation with higher concentrations of the inhibitor led to a significant inhibition in β-arrestin 2 recruitment when compared to buffer incubated cells. Values are given as mean +/− SEM (n=9 of three independent experiments); *p<0.05.

β-arrestin 2 recruitment depends at least in part on receptor phosphorylation by GPCR kinases (GRKs). To test whether this dependency can be demonstrated by the A_1_AR NanoBit^®^ assay, cells were preincubated with a specific inhibitor of GRK2 (βARK1 inhibitor). Data presented in figure 5 B demonstrate the dose dependent inhibition of β-arrestin 2 recruitment through incubation with GRK2 inhibitor which was statistically significant at 100 μM.

### Valerian extract Ze 911 induces A_1_AR mediated β-arrestin 2 recruitment

A_1_AR plays an important role in the regulation of sleep. In this context, valerian extracts have been demonstrated to show agonistic activity upon A_1_AR, maybe explaining the sleep-inducing effect of this phytopharmaceutical. Until today no data regarding a possible influence of valerian on the β-arrestin 2 recruitment via A_1_AR exists. As demonstrated by data in figure 6 A valerian extract Ze 911 induced a robust A_1_AR mediated recruitment of β-arrestin 2. EC_50_ of this activation was calculated to be 78 μg/ml.

**Figure 6:**
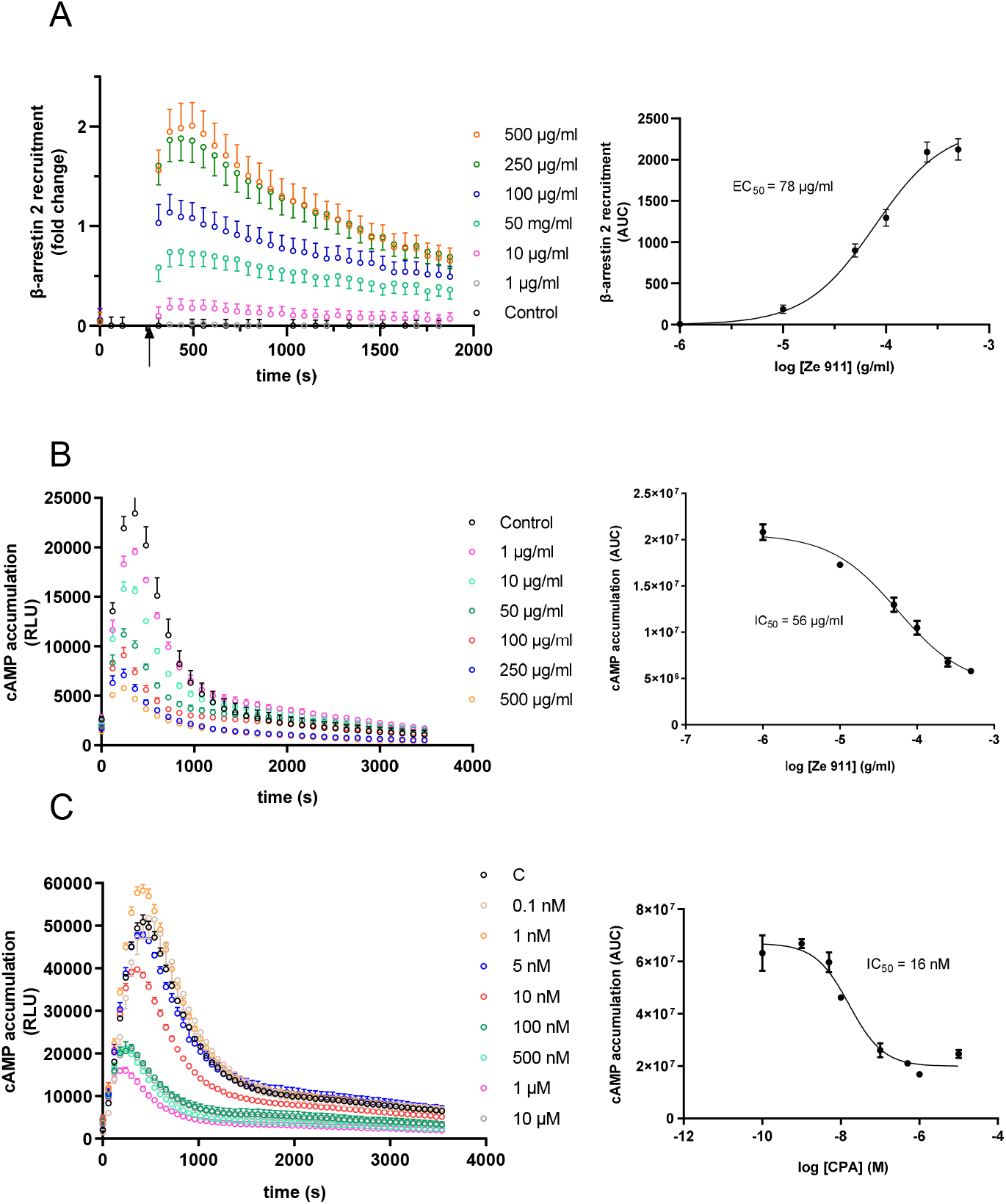
(A) Concentration-dependency of β-arrestin 2 recruitment in the A_1_AR Nano-Bit^®^ reporter assay: At the time point indicated by the arrow valerian extract Ze 911 was added to A_1_AR-Nano-Bit^®^-βarr2 HEK 293 cells at concentrations indicated beside the graph and luminescence was measured for 30 minutes. A solvent control of 0.1% DMSO was included. Dose response curves on the right were calculated using the areas under the curves (AUC). Values are given as mean +/− SEM (n=9 of three independent experiments). Concentration-dependency of Gi activation measured by the inhibition of cAMP production in the GloSensor^TM^ reporter assay for (B) valerian extract Ze 911 and (C) CPA: HEK GloSensor^®^ cells were incubated with concentrations indicated beside the graph and cAMP accumulation was stimulated with 1 μM isoprenaline + 1 μM forskolin (timepoint 0) and luminescence was recorded for 60 minutes. A solvent control of 0.1% DMSO was included. Dose response curves on the right were calculated using the area under the curves (AUC). Values (mean ± SEM) of three wells from an exemplary experiment are shown. Experiments were repeated with similar results (see SI Figure 2).

Since the literature focused on G_αi_ activation by valerian and not β-arrestin recruitment a cAMP assay was performed to compare the results (Fig. 6 B). A_1_AR overexpressing HEK GloSensor™ cells were used to measure the influence of valerian extract on cAMP accumulation after β-adrenergic stimulation with isoprenaline. Cells were treated with different concentrations of valerian extract. The concentration of cAMP was directly measured after treatment. The cAMP concentration decreased with increase of isoprenaline concentration. The IC_50_ was calculated to be 56 μg/ml. For comparison, cAMP data for CPA were collected in the same manner as for valerian extract (Fig. 6 C; for replicates see SI Fig. 2 B).

## Discussion

Originally identified as adapter proteins mediating receptor desensitization and internalization, β-arrestins are now a recognized component of GPCR signal transduction (Chen and Tesmer, 2022; Jiang et al., 2022). While for some GPCRs the interaction with β-arrestins is well studied, there is little data on the recruitment of these proteins by A_1_AR. Therefore, in this work, an assay based on NanoBit^®^ technology was established and examined in more detail to determine how well it is suited for studying the activation of A_1_AR and subsequent recruitment of β-arrestin 2 (Dijon et al., 2021).

Using the newly established NanoBit^®^ assay, we determined an EC_50_ value of 884 nM for adenosine for A_1_AR mediated β-arrestin 2 recruitment. In comparison, IC_50_ values for adenosine ranging from of 100 to 310 nM were found for adenylate cyclase mediated cAMP formation (Fredholm et al., 2001; Müller & Jacobson, 2011; Yan et al., 2003). These differences can be explained by differences in the assays themselves. For example, whereas A_1_AR-cAMP assays indirectly measure the inhibitory effect of receptor activation on adenylate cyclase activity stimulated by another substance such as forskolin, the assay presented here directly measures the activity of A_1_AR via immediate β-arrestin 2 recruitment. It has been reported that receptor density affects values of A_1_AR activities obtained by the same agonist (Cordeaux et al., 2000). In addition, concentrations of endogenously produced adenosine or inosine, which may be different in cell cultures and cell membrane preparations, also affect the A_1_AR activity found in the different assays (Cohen et al., 1996; Jarvis and Thompson, 2019).

Nevertheless, the values determined with the different assays are in the same medium to high nanomolar range and are therefore comparable.

For CPA, Mueller et al. determined an IC_50_ value of 24 nM for inhibition of cAMP production (Müller et al., 2002), which is about five to six times lower than what was found here for β-arrestin 2 recruitment (129 nM). Again, the assays are only partially comparable because cAMP accumulation was measured on isolated membranes of A_1_AR-overexpressing CHO cells treated with adenosine deaminase (ADA), whereas our NanoBit^®^ assay was performed on live HEK cells that were not pretreated with ADA. Therefore, we additionally performed a cAMP assay on live HEK cells (GloSensor™ assay). The IC_50_ value of 16 nM for CPA that we obtained in this assay fits perfectly with the results of Mueller and coworkers (Müller et al., 2002). Interestingly, the five to sevenfold ratio between cAMP inhibition and β-arrestin 2 recruitment for CPA corresponds to that described above for adenosine (100 - 310 nM to 884 nM). For the nonspecific AR agonist NECA, an EC_50_ value of 115 nM for β-arrestin-2 recruitment was found to be similar to that for CPA. In contrast, the IC_50_ value of 56 nM determined for NECA on A_1_AR by Alnouri et al. for cAMP accumulation is slightly higher than the value determined by the same research group for CPA (Alnouri et al., 2015). This poorer potency of NECA compared to CPA regarding the inhibition of cAMP production via A_1_AR is also confirmed by studies of Cordeaux and coworkers (Cordeaux et al., 2000).

Recruitment of β-arrestins has also become a focus of scientific investigation in recent years because of the tremendous increase in our understanding of the importance of the interplay between the various signaling pathways of GPCRs in the development of diseases, as well as in the efficacy or side effect profile of drugs (Sharma and Parameswaran, 2015; Bagnato and Rosanò, 2019; Bond et al., 2019; Zhai et al., 2022). There are already first drugs that exploit the effect of preferred activation of one of several possible signal transduction pathways also called biased signaling, such as the cardiac drug carvedilol (Wisler 2007; Ibrahim 2012; Kolb 2022; Kenakin 2019). A positive effect of biased agonism on the side effect profile is also thought to be present for A_1_AR (Valant 2014; Baltos 2016; McNeill 2021). For example, benzyloxy-cyclopentyladenosine has been identified as a selective A_1_AR agonist that achieves analgesia without the adverse side effect of cardiorespiratory depression (Wall et al., 2022).

As a non-nucleoside and biased agonist, we chose capadenoson which was developed by Bayer as a cardioprotective, highly selective A_1_AR agonist with an improved safety profile (Albrecht-Küpper et al., 2012; Sabbah et al., 2013). To our knowledge there is no data regarding β-arrestin 2 recruitment mediated by capadenoson activated A_1_AR. The established NanoBit^®^ assay detected a pronounced partial agonism for this compound, whereas it showed full agonism in case of adenylate cyclase inhibition. This fits to data collected by Baltos et al. who demonstrated a slight increase in cAMP inhibition as well as a reduction of ERK1/2 and Akt phosphorylation after capadenoson stimulation compared to NECA (Baltos et al., 2016). ERK1/2 phosphorylation in response to A_1_AR activation is at least partially β-arrestin-dependent (Jajoo et al., 2010). Therefore, the reduced ERK1/2 phosphorylation as described by Baltos et al. in the case of capadenoson can be explained by the reduced recruitment of β-arrestin 2 demonstrated in our work.

To verify the assay as a tool for bias calculation, we measured all agonists in saturated conditions in both the Glosensor^TM^ cAMP as well as the NanoBit^®^ assay and compared their activities based on calculation of the AUCs. In this way a pronounced bias for A_1_AR mediated G protein activation compared to β-arrestin 2 recruitment was prooven for capadenoson. While capadenoson acts as a full agonist on the cAMP level it only acts as a partial agonist for β-arrestin 2 recruitment. Both NECA and adenosine had a tendency to activate recruitment of β-arrestin 2 more than lowering cAMP levels compared with CPA, which fits the data discussed above (Cordeaux et al., 2000; Alnouri et al., 2015). On the other hand, it has to be taken into account that the unspecific agonists NECA and adenosine activate other ARs like A_2A_AR and A_2B_AR thereby increasing the cAMP level and distorting the cAMP reduction mediated by A_1_AR. This point is neglectable in case of the β-arrestin 2 assay because of its specificity for A_1_AR. Therefore, the calculated shift to β-arrestin 2 recruitment calculated for NECA and adenosine might be overestimated. A more specific assay addressing the inhibition of cAMP production like a Gαi protein recruitment assay comparable to the NanoBit^®^ assay for β-arrestin 2 would solve this uncertainty (Nehmé et al., 2017; Höring et al., 2020; Laschet and Hanson, 2021).

When calculated via AUCs, A_1_AR mediated β-arrestin 2 recruitment was a 2.2 and 2.4-fold higher for adenosine, CPA and NECA compared to capadenoson. It should be pointed out that this calculation includes not only the activation of the receptor, but also its deactivation/internalization. However, this is also significantly lower in the case of capadenoson due to its reduced activation behavior. To compare different GPCR agonists it can therefore be useful to characterize the activation phase of the receptor only. Hoare and coworkers developed an alternative tool to analyze kinetic signaling data from fluorescent reporter systems ((Hoare et al., 2021; Hoare, Tewson, Quinn, & Hughes, 2020; Hoare, Tewson, Quinn, Hughes, et al., 2020); available as plug ins for Prism). These plug-ins calculated curves that matched our measurement data very well. Initial rates that reflect the activation kinetics of the different agonists can be calculated from these curves. The difference calculated on the basis of the initial rates (up to 8.8-fold) was significantly higher than the difference calculated on the basis of AUCs (see above), more clearly reflecting different receptor activation by the agonists tested here. Calculating the initial rate is therefore probably the more sensitive way to investigate different agonists in case of β-arrestin 2 recruitment measured by the NanoBit^®^ assay.

Assay systems for G protein activation and arrestin recruitment are of interest for agonist screening but also for testing receptor modulators that might shift signaling to one pathway or the other. In this context, the results for VCP 171 are remarkable because they show that the assay can be readily adapted to determine the activity of modulating agents. The increase in agonistic effect of NECA between 10 and 20% fits data reported in the literature (Aurelio et al., 2009; Nguyen et al., 2016). Inhibitory substances for the A_1_AR can also be screened without problems, as shown here exemplarily for DPCPX. The IC_50_ of 124 nM that we calculated fits well with values measured on a model for depression of synaptic transmission mediated by the A_1_AR for DPCPX (Latini et al., 1999).

Internalization of A_1_AR is a slow process compared to A_3_AR (Klaasse et al., 2008). It is known that β-arrestin recruitment is regulated in part through receptor phosphorylation by GPCR kinases (GRKs), but phosphorylation of A_1_AR by GRKs is still controversial (Klaasse et al., 2008; Nie et al., 1997; Ramkumar et al., 1993; Soave et al., 2020). However, in the assay described here, inhibition of GRK2-mediated receptor phosphorylation results in decreased β-arrestin 2 recruitment. The IC_50_ value of 126 μM published by Iino et al. for the βARK-1 inhibitor fits our observations that efficient inhibition of β-arrestin 2 recruitment starts at 50 μM and higher (Iino et al., 2002). Therefore, A_1_AR mediated β-arrestin 2 recruitment is at least in part regulated by GRK phosphorylation of the receptor.

After testing chemically defined substances for activating the A_1_AR we decided to investigate a more complex compound mixture with agonistic activity. Phytopharmaceuticals containing valerian extracts are used as mild sleep-inducing agents (for review see (Borrás et al., 2021; Shinjyo et al., 2020)). There is evidence in the literature that this effect is at least partly dependent on A_1_AR activation (Lacher et al., 2007; Müller et al., 2002; Schumacher et al., 2002; Vissiennon et al., 2006). All data on a possible influence of valerian extract or its ingredients on A_1_AR are based on receptor binding data as well as cAMP accumulation. Data on the regulation of β-arrestin 2 recruitment by A_1_AR activation mediated by valerian extracts have not yet been published. The half maximal inhibition of cAMP accumulation for the valerian extract used in our study was reached at 56 μg/ml in the GloSensor™ assay, whereas it was 900 μg/ml in the activated charcoal absorption assay of Müller et al. (Müller et al., 2002). In addition to the differences in the cAMP assays used, the composition of the valerian extract tested by Mueller and coworkers is not directly comparable to Ze 911 investigated here, which explains the different IC_50_ values. However, despite all differences, comparable inhibition of cAMP accumulation is described in both publications. Interestingly, β-arrestin 2 recruitment is only 1.5-fold higher than cAMP inhibition for Ze 911, with an EC_50_ value of 78 μg/ml, and thus significantly lower than the five to sevenfold ratio described for CPA above. Ze 911 therefore appears to have a slight bias toward the recruitment of β-arrestin 2 compared to adenosine or CPA.

The NanoBit^®^ assay for β-arrestin 2 recruitment has been previously reported as a useful tool for other adenosine receptors (Storme, Cannaert, et al., 2018; Storme, Tosh, et al., 2018). Alternatively, arrestin recruitment has been measured by BRET, FRET or complementation of β-galactosidase (Gao & Jacobson, 2008; Navarro et al., 2020). Compared to FRET/BRET, the signal-to-noise ratio of Nano-Bit^®^ assays is very high. For example, signal increases by a factor of 10 can be clearly observed in our assay. This makes the assay extremely sensitive and may allow the establishment of recruitment assays with physiological expression levels of receptors and arrestins in the future. This possibility is also favored by the small size of 19 kDa of the complemented enzyme (Wouters et al., 2018). Initial approaches to introduce the NanoBit^®^ system into cells at the genomic level already exist (Dale et al., 2019; Oh-hashi et al., 2017; White et al., 2019).

In conclusion, the presented assay is very well suited to study A_1_AR mediated recruitment of β-arrestin 2 by different substances. Together with other assays like the cAMP GloSensor™ assay used here and appropriate tools for evaluation, the analysis of biased signaling is also very feasible with this assay.

## Authors Contribution

LS performed the experiments. LS and SF wrote the manuscript. HH and SF designed the experiments and acquired funding for the study. HH edited the manuscript.

## Acknowledgements

We thank Samuel R. J. Hoare from Pharmechanics LLC, Owego, New York for providing the plug ins for GraphPad Prism and for helpful discussions about our results.

## Supplemental Information

**SI Figure 1:**
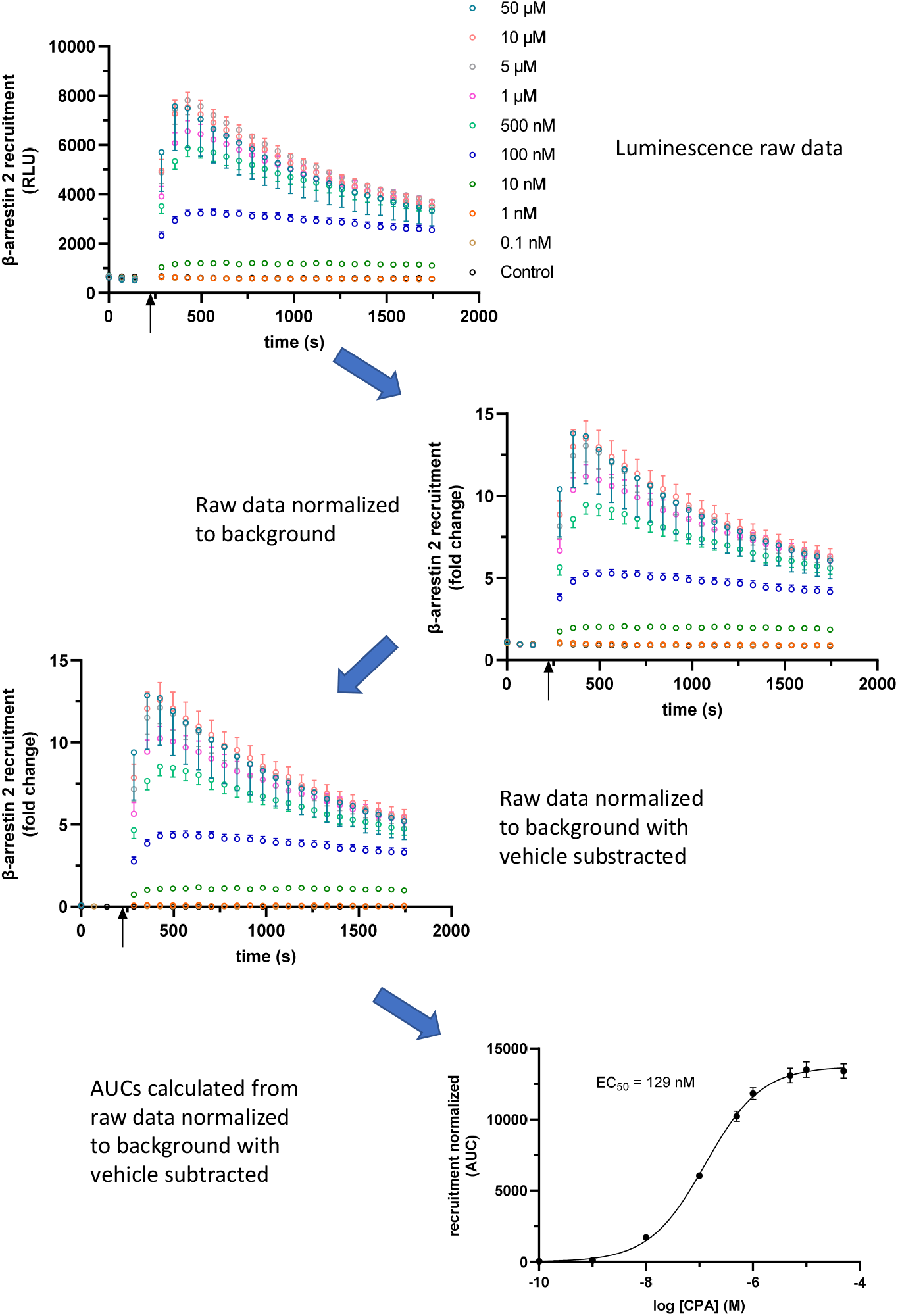
Processing of raw data. A_1_AR NanoBit^®^ reporter assay: At the time point indicated by the arrow CPA was added to A_1_AR-NanoBit^®^-βarr2 HEK 293 cells at concentrations indicated beside the graph and luminescence was measured for 30 minutes. A solvent control of 0.1% DMSO was included. Data for relative luminescence light units (RLU) were transferred to Prism and plotted as a function of time. To normalize for well to well variabilities each data point was divided by the mean of the first three values using the “Remove baseline and column math” function of the software. Next the solvent control was subtracted using the same functionality of Prism. From this data set areas under the curves were calculated using the corresponding function and plotted against the concentration of the agonist used (log [agonist] (M)) to receive a dose/response dependency. For each value RLUs or AUCs (mean ± SEM) of nine wells from three different experiment are shown.

**SI Figure 2:**
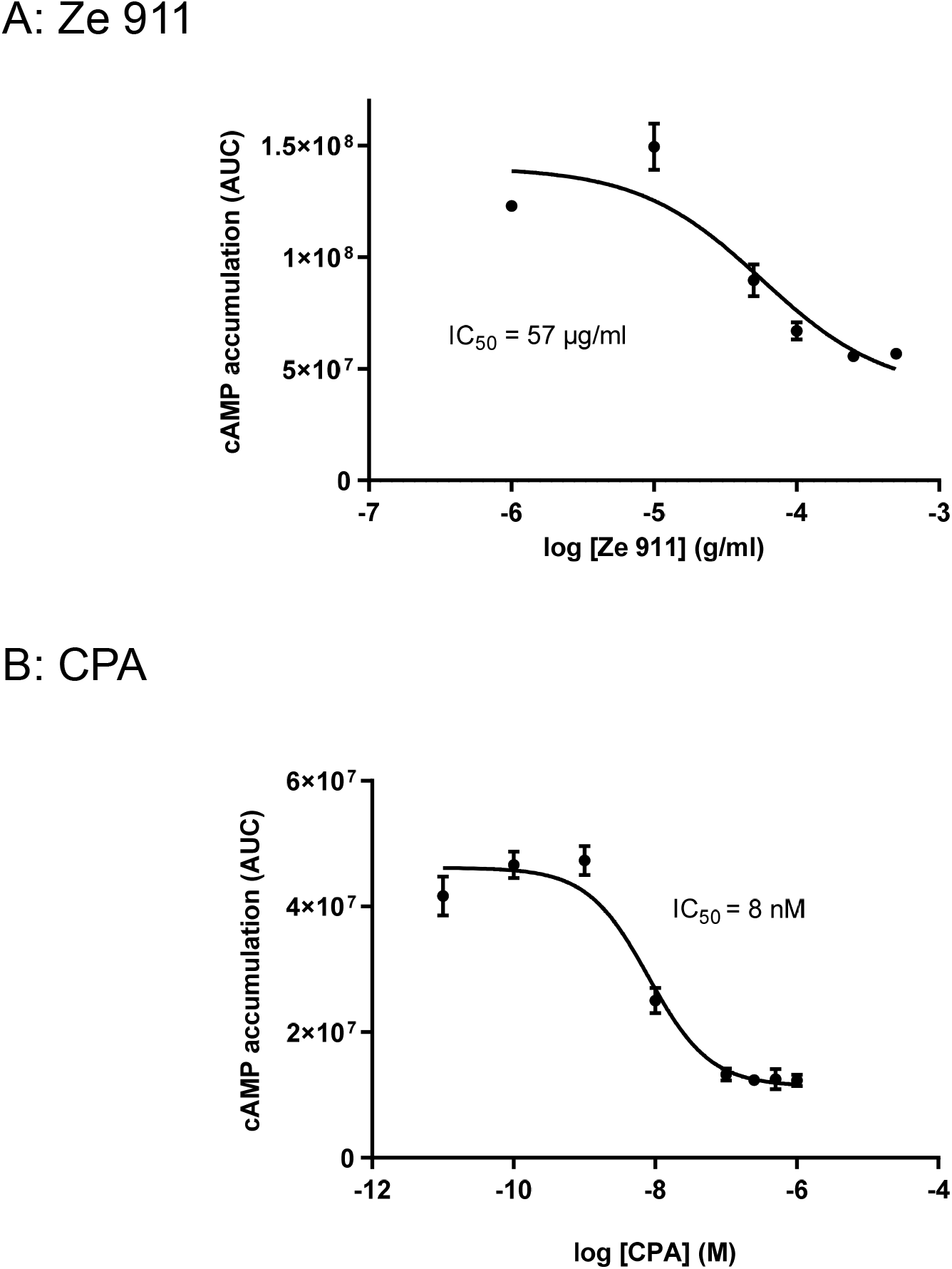
Replicates of cAMP experiments. Concentration-dependency of Gi activation measured by the inhibition of cAMP production in the GloSensor™ reporter assay for (A) valerian extract Ze 911 and (B) CPA: HEK GloSensor^TM^ cells were preincubated with concentrations indicated and cAMP accumulation was stimulated with 1 μM isoprenaline + 1 μM forskolin (timepoint 0) and luminescence was recorded for 60 minutes. A solvent control of 0.1% DMSO was included. Dose response curves calculated using the area under the curves (AUC). Values (mean ± SEM) of three wells are shown. See also regular Figure 6.

